# SCANVIS – a tool for SCoring, ANnotating and VISualizing splice junctions

**DOI:** 10.1101/549261

**Authors:** Phaedra Agius, Heather Geiger, Nicolas Robine

## Abstract

**Motivation:** The association of splicing signatures with disease is a leading area of study for prognosis, diagnosis and therapy, frequently requiring detailed analysis of splicing events across multiple samples. We present a novel fast-performing annotation-dependent tool called SCANVIS for scoring and annotating splice junctions by gene name, junction type and any frame shifts incurred. SCANVIS has a novel and fast visualization technique that distinguishes annotated splice junctions from unannotated ones in the context of nearby variants and read coverage. It also allows users to merge samples across cohorts, thereby allowing for quick comparisons of splice junctions across diseases and tissue types.

**Results:** We show that SCANVIS generates reasonable PSI scores by demonstrating that tissue/cancer types in GTEX and TCGA are well separated and easily predicted from a few thousand SJs. We also show how SCANVIS can be used to map out junctions overlaid with variants and read coverage for one or more samples, with line types and colors delineating frame shifts and junction types.

**Availability:** SCANVIS is available for download at https://github.com/nygenome/SCANVIS

## 1 Introduction

The association of splicing defects with the occurrence of disease and variants is an ongoing area of research. Some splicing analysis tools operate independently from annotation [1,2] making it difficult for users to quickly identify genes harboring aberrant splice junctions (SJs) relevant to the disease. Frequently tools lack the details crucial to inferring potential downstream consequences of a SJ. For instance in-frame SJs supported by annotation are less likely to be aberrant than frame-shifting un-annotated SJs. It is also useful to be able to visualize SJs and nearby mutations. To that end, IGV sashimi plots are useful but laborious and memory intensive when viewing multiple samples, and it is difficult to distinguish annotated from unannotated SJs and frame-shift details. We developed SCANVIS in an effort to address these short-comings. It is an R-based tool that scores and annotates SJs with several details such as gene name, junction type and frame-shifts. It has a visualization component that quickly generates sashimi-style plots for one or more samples for a given genomic region, optionally overlaid with variants and read coverage. SCANVIS also has a merge function that allows users to average SJ signatures and count variants across multiple samples in a cohort, thereby allowing users to quickly glimpse the cohort’s SJ landscape in one figure.

## 2 Methods

SCANVIS requires two main inputs to process a sample. The first is a set of SJ coordinates with read support – these details can be derived from an alignment algorithm of choice. In our analyses we used the SJ files generated by the STAR aligner [Dobin *et al*]. Different software packages may represent SJ coordinates in different ways – both STAR and SCANVIS assume the start/end position of an annotated splice junction (ASJ) to be the intronic position adjacent to the exons. The second required input is an annotation file with details derived from a gtf file using an inbuilt function in our software. In our analyses we used human gen-code 19 which is supplied with the software package.

By design SCANVIS scores and annotates one sample at a time with PSI (Percent Spliced-In) scores defined as ratios of SJ reads and computed at local intervals, thereby operating within the distribution of SJ reads for that sample. This individualistic and localized approach reduces batch effects and deters a poor quality sample from influencing the rest of the cohort. For each junction in a sample, SCANVIS delivers the following details: *(i)* genomic interval used for PSI-scoring, *(ii)* PSI score, *(iii)* names of genes overlapping the SJ, *(iv)* junction type with unannotated SJs (USJs) described as one of the following: exon-skip, alt5p, alt3p, IsoSwitch, Unknown, Novel Exon (NE); and *(v)* any frame-shifts, whether the SJ is in frame across all isoforms, or not, and if mixed then each isoform and frame consequence is listed. The definitions of exon-skip, alt5p and alt3p are as typically described in splicing literature. IsoSwitch events are defined to be SJs that straddle two mutually exclusive isoforms and Unknown events correspond to SJs that have start and end coordinates landing in intronic regions. Novel Exons are described in detail below.

The PSI (Percent Spliced-In) score is defined to be the ratio of the number of reads *x* supporting a query junction to the sum of *x* and *y*, where *y* is the expected number of ASJ reads in a genomic interval containing the query junction. Genomic intervals are constructed by merging gene and SJ coordinates so that overlapping genomic regions are condensed into one region, provided the following two constraints are met: every SJ must be contained in exactly one interval and an interval must contain at least one ASJ. Then *y* is defined to be the median of all ASJ reads in the interval. This approach spares the PSI denominator from contamination by spurious unannotated SJs (USJs). Since USJs frequently have poor read support and may be alignment artifacts, it is worth reducing their influence on the PSI computation.

In addition to the user-supplied SJs, SCANVIS also lists the start/end coordinates of all USJ pairs coinciding in intronic regions, and describes these as Novel Exons (NE). These are scored by the mean PSI of all SJs landing on the start/end coordinates. If the bam file is supplied, a second read-coverage based PSI is also computed. This is defined as *c*/(*c5*+*c*+*c3*) where *c* is the mean read coverage of the NE, and *c5* and *c3* are the mean read coverages for flanking regions, both defined to be intervals of length 20% of the NE interval. Running the tool with the bam file takes longer but is recommended for more thorough NE support.

## 3 Results

We ran SCANVIS on thousands of samples from TCGA [http://can-cergenome.nih.gov/] and GTEX [Carithers *et al*] which we downloaded and aligned to hg19 using the STAR aligner [Dobin *et al*]. We explored 3706 TCGA samples across 9 cancer types and 4082 GTEX samples over 14 tissue types. Each tissue type averaged 338 samples with TCGA types BRCA and HNSC being the largest and smallest groups (1220 and 24 samples respectively, see Supplementary Figure S1). As a first assessment of the SCANVIS PSI scoring system, we explored tissue differentiation using t-SNE plots computed on the top 5000 most variable SJs. In Figure 1(a,b) we observe good tissue separation for GTEX and TCGA samples. We also computed a joint t-SNE plot on both datasets using 1528 SJs in the intersection of the top 5000 SJs for TCGA and GTEX. Figure1(c) shows that tissue separation precedes database separation, suggesting that the inherent normalization in the PSI scoring system is reasonably resistant to batch effects. We further explored tissue separation by predicting tissue types, using linear kernel SVMS on the PSI scores in a 10-fold cross-validation learning system over 20 iterations. At each learning iteration, we selected the top 500 most variable SJs from the top 5000 SJs used for the t-SNE plots, with variance computed on the training samples only. In Supplementary Figure S2 we show that the resulting AUC scores (area under ROC plots) exceed 0.98 for most tissue and cancer types.

**Figure 1.**
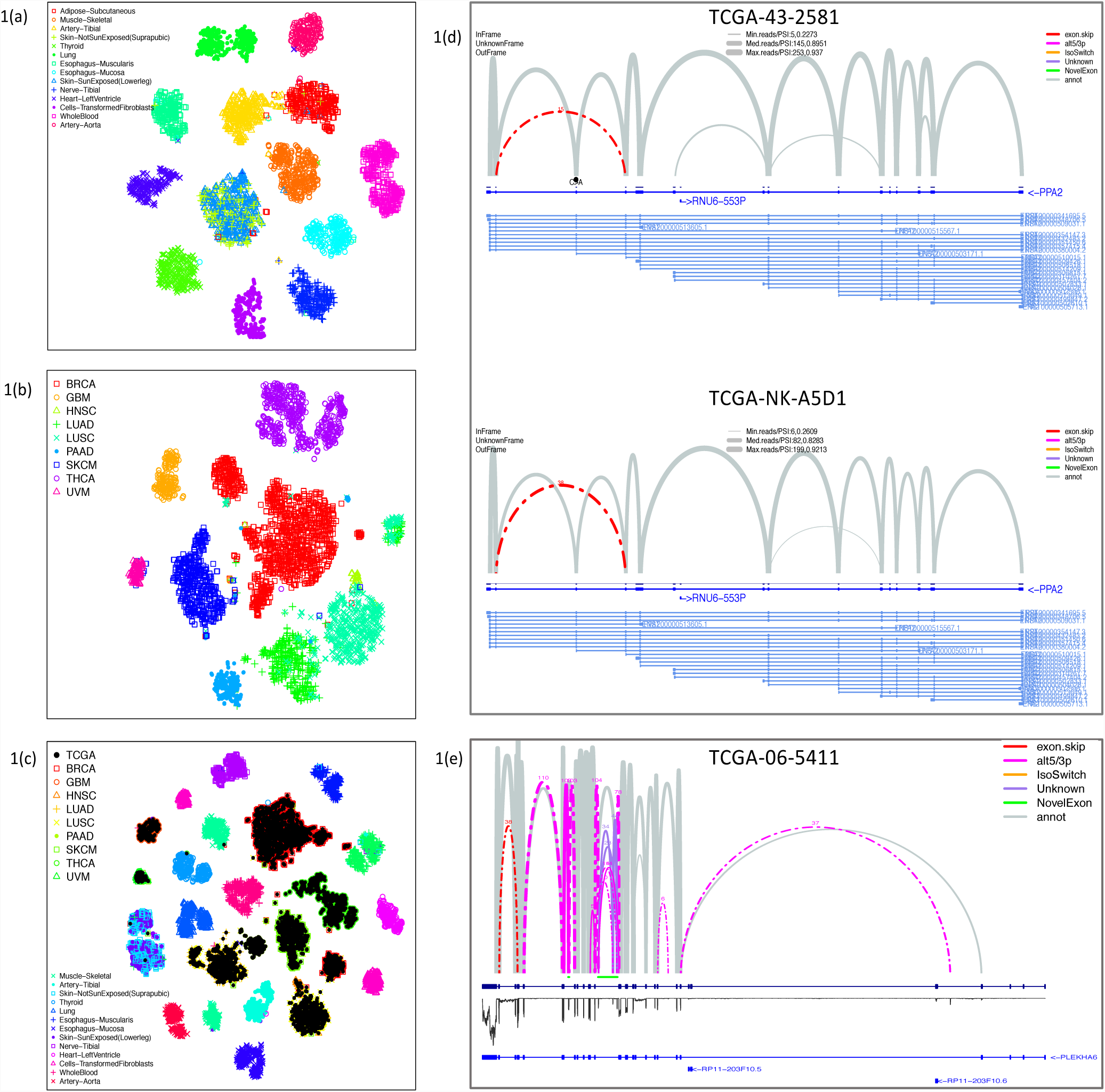
t-SNE plots (left) and SCANVIS visualization plots (right) The t-SNE plots in (a,b) were computed using SCANVIS PSI scores on the top 5000 most variable SJs in our GTEX and TCGA cohorts, while (c) was computed on 1528 SJs that were in the intersection of top 5000 SJs for GTEX and TCGA. Figure 1(d) shows splice junctions in the pyrophosphatase gene PPA2 for two LUSC samples from TCGA, and an exon skipping event that was exclusive to these two samples. The top LUSC sample harbors a variant in the splice region associated with the exon being skipped, suggesting that the variant could be having some impact for this sample. Figure 1 (e) shows a SCANVIS plot of the pleckstrin homology domain (PLEKHA6), with splicing signatures overlaid with the read coverage for a GBM sample. A number of USJs coincide in the intronic regions (marked in green). The three small peaks in the read coverage in this region suggest there is some read support for Novel Exons in this area.

The visualization tool is arguably the most useful SCANVIS component as it allows users to quickly project SJs in sashimi-style plots together with variants and read coverage in one or more samples. The resulting figure overlays ASJs in grey arcs with USJs in various colors indicating USJ types (exon skipping, alt5/3p etc). SJ arc heights relate to the number of reads and line widths correlate with SCANVIS PSIs, while line types correspond to in-frame and out-of-frame junctions. SCANVIS includes an optional function to map variants to SJs by overlapping their coordinates. If variant-mapped SJs are submitted to the visualization tool, the tool generates a figure overlaying variants with SJs, with single point mutations shown as black dots and structural variants as black lines. Figure 1d (top) shows an exon skipping event (red arc) in the gene PPA2 that was found in 2 LUSC samples and no other TCGA samples, with one of those samples having a nearby variant in a splice region. Overlaying the read profile can be quite useful for visualizing Novel Exons and for understanding local support splice junctions. When users submit the bam file associated with an individual sample, our visualization tool delivers a sashimi plot overlaid with a read coverage profile. Figure 1e shows SJs in the gene PLEKHA6 for a GBM sample, with the read coverage shown as an inverted black profile. A region ripe with alternative 5p and 3p events is marked in green as a Novel Exon in this example, and the read profile indicates support for at least two significant peaks in that area.

One last additional feature of the visualization tool is that multiple samples can be merged to produce one sashimi figure, allowing for users to quickly contrast sample cohorts (eg disease versus control samples). When samples are merged, the union of all SJs is collected over the samples, and the PSI and number of supporting reads are averaged across the samples. If variant-mapped SJs are submitted, the mutations are also counted, and are collectively represented as a lollipop plot when the same mutation is observed in more than one sample.

## 4 Discussion

Being able to determine which genes harbor splicing defects and which splicing defects incur frame shifts are critical questions in splicing analyses. SCANVIS scores and annotates SJs with attention to many details that can assist in the inference of any downstream consequences. It runs quite efficiently - a 1.6G MacBook Air with 8G memory processed a sample with 170K SJs (without the bam file supplied) in 6 minutes. Its efficiency in generating multiple sashimi-style plots is also worth emphasizing. We used this tool extensively for the analysis of in-house samples, and we hope that in making this tool accessible to the community, researchers can dig deeper into splicing signatures with more transparency.

## Acknowledgements

We thank Christian Stolte for valuable suggestions on how to make SCANVIS sashimi visualizations look better.

## References

Yang I. Lee et al (2018) Annotation-free quantification of RNA splicing using Leaf-Cutter. Nature Genetics 50, 151–158.

Qingqing Wang and Donald C. Rio (2018) JUM is a computational method for comprehensive annotation-free analysis of alternative pre-mRNA splicing. PNAS, 115(35).

Alexander Dobin et al (2013) STAR: ultrafast universal RNA-seq aligner. Bioinformatics, 29(1) 15–21.

Carithers, Latarsha J, Ardlie, Kristin, Barcus, Mary, Branton, Philip A, Britton, Angela, Buia, Stephen A, Compton, Carolyn C, DeLuca, David S, Peter-Demchok, Joanne, Gelfand, Ellen T, Guan, Ping, Korzeniewski, Greg E, Lockhart, Nicole C, Rabiner, Chana A, Rao, Abhi K, Robinson, Karna L, Roche, Nancy V, Sawyer, Sherilyn J, Segrè, Ayellet V, Shive, Charles E, Smith, Anna M, Sobin, Leslie H, Undale, Anita H, Valentino, Kimberly M, Vaught, Jim, Young, Taylor R, Moore, Helen M, on behalf of the GTEx consortium (2015) A Novel Approach to High-Quality Postmortem Tissue Procurement: The GTEx Project, Biopreservation and Biobanking 13(5), 2015, p 311–319

