## Supplemental Figures S1 and S2 for "SCANVIS – a tool for SCoring, ANnotating and VISualizing splice junctions"

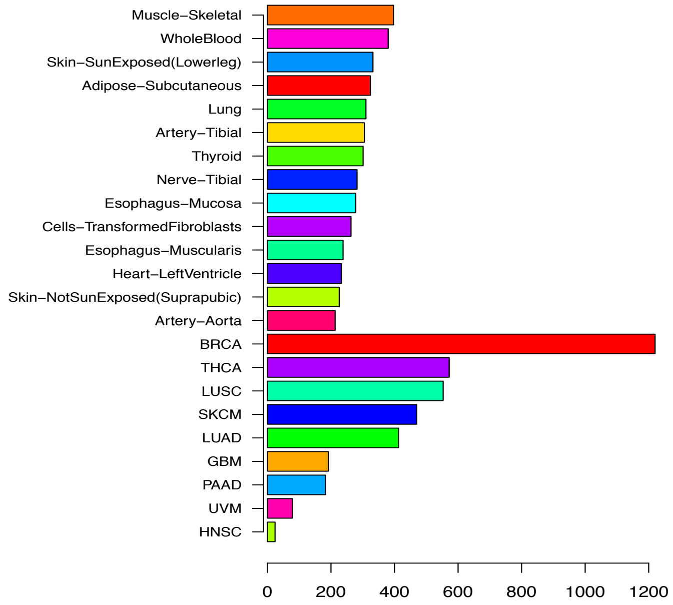


**Figure S1. Sample count barplots.** Each bar shows the number of samples per cancer type in the set of TCGA samples that we used (bottom 9 bars) and the number of samples per tissue type in GTEX (top Artery-Aorta to Muscle-Skeletal)

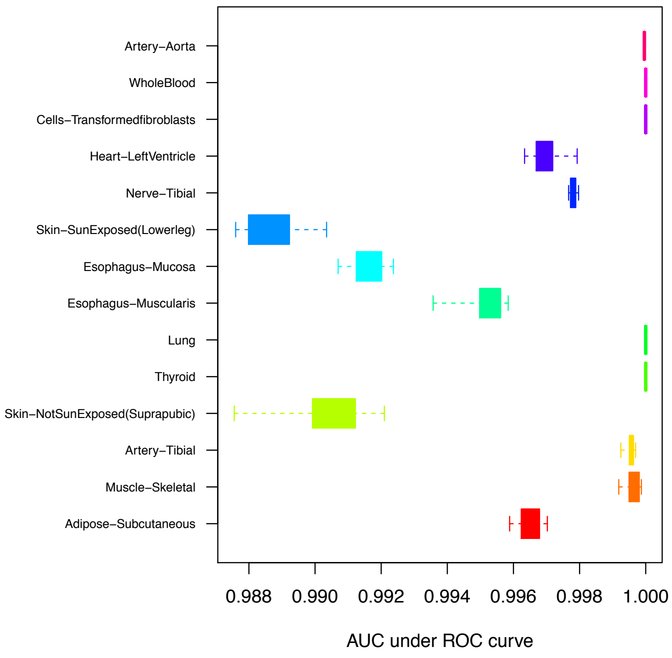


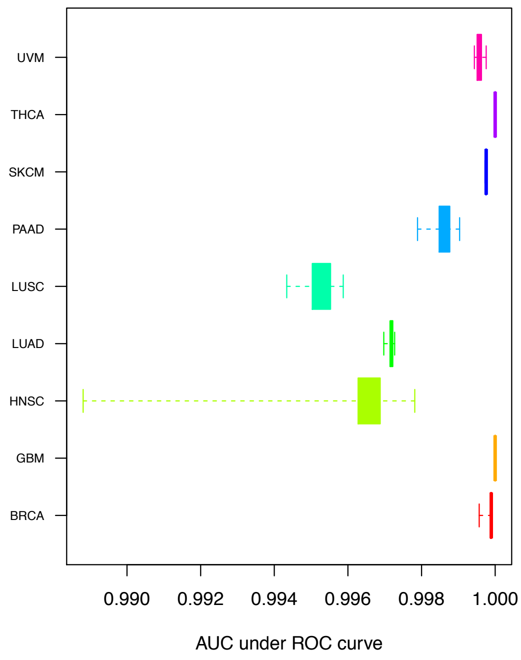


**Figure S2. AUC barplots** These figures show AUC boxplots over the 20 iterations of 10-fold cross-validation runs predicting whether samples belong to a certain tissue/cancer type or not (left figure for GTEX, right figure for TCGA samples). The top 500 most variable SJs were selected as features at each cross-fold run, with variance computed on the training samples, and we used a linear SVC (Support Vector Classification) as our modeling system. Most tissue types and cancer types were easily predicted with AUCs above 0.98. However we observed a slight drop in AUC scores for skin tissues in the GTEX cohort, presumably because sun-exposed skin is quite a bit similar to unexposed skin than it is to other tissue types in the cohort. We also observe a relatively wide range of AUCs for the cancer type HNSC – this is explained by the small number of samples comprising this group (just 24) making it harder for the modeling system to pick up SJs relative to HNSC during training.
